# FtsEX-mediated regulation of inner membrane fusion and cell separation reveals morphogenetic plasticity in *Caulobacter crescentus*

**DOI:** 10.1101/124214

**Authors:** Elizabeth L. Meier, Qing Yao, Allison K. Daitch, Grant J. Jensen, Erin D. Goley

**Affiliations:** Department of Biological Chemistry, Johns Hopkins University School of Medicine, Baltimore, Maryland 21205, USA; Division of Biology and Biological Engineering, Pasadena, California 91125, USA; Howard Hughes Medical Institute, California Institute of Technology, Pasadena, California 91125, USA

**Author notes:** Correspondence and requests for materials should be addressed to E.D.G.

## Abstract

During its life cycle, *Caulobacter crescentus* undergoes a series of coordinated shape changes, including generation of a polar stalk and reshaping of the cell envelope to produce new daughter cells through the process of cytokinesis. The mechanisms by which these morphogenetic processes are coordinated in time and space remain largely unknown. Here we demonstrate that the conserved division complex FtsEX controls both the early and late stages of cytokinesis in *C. crescentus*, namely initiation of constriction and final cell separation. Δ*ftsE* cells display a striking phenotype: cells are chained, with skinny connections between cell bodies resulting from defects in inner membrane fusion and cell separation. Surprisingly, the thin connections in Δ*ftsE* cells share morphological and molecular features with *C. crescentus* stalks. Our data uncover unanticipated morphogenetic plasticity in *C. crescentus*, with loss of FtsE causing a stalk-like program to take over at failed division sites and yield novel cell morphology.

**Author Summary:** Bacterial cell shape is genetically hardwired and is critical for fitness and, in certain cases, pathogenesis. In most bacteria, a semi-rigid structure called the cell wall surrounds the inner membrane, offering protection against cell lysis while simultaneously maintaining cell shape. A highly dynamic macromolecular structure, the cell wall undergoes extensive remodeling as bacterial cells grow and divide. We demonstrate that a broadly conserved cell division complex, FtsEX, relays signals from the cytoplasm to the cell wall to regulate key developmental shape changes in the α-proteobacterium *Caulobacter crescentus*. Consistent with studies in diverse bacteria, we observe strong synthetic interactions between *ftsE* and cell wall hydrolytic factors, suggesting that regulation of cell wall remodeling is a conserved function of FtsEX. Loss of FtsE causes morphological defects associated with both the early and late stages of division. Intriguingly, without FtsE, cells frequently fail to separate and instead elaborate a thin, tubular structure between cell bodies, a growth mode observed in other α-proteobacteria. Overall, our results highlight the plasticity of bacterial cell shape and demonstrate how altering the activity of one morphogenetic program can produce diverse morphologies resembling those of other bacteria in nature.

## Introduction

Bacteria are capable of adopting an impressive array of shapes exquisitely tuned for their particular environmental niches. Underpinning these shapes is the bacterial cell wall, which plays an essential role in specifying and maintaining diverse morphologies [1]. The cell wall consists of a layer of peptidoglycan (PG) composed of glycan strands of repeating disaccharide subunits crosslinked by pentapeptide bridges. In addition to adapting to changing environments, the PG also undergoes dynamic remodeling to drive shape changes during dedicated cellular processes such as division [2,3].

The α-proteobacterium *Caulobacter crescentus* is an ideal model organism for the study of cell shape as it undergoes a series of coordinated morphogenetic changes during its cell cycle. After every division event, *C. crescentus* produces two distinct daughter cell types. One is a flagellated, motile swarmer cell, which contains a flagellum and pili at one cell pole [4]. The other is a sessile stalked cell, where the polar flagellum has been replaced by a thin, tubular extension of the cell envelope known as a stalk [4]. Unable to replicate its chromosome or initiate cell division, a swarmer cell differentiates into a stalked cell by ejecting its flagellum, disassembling its pili, and growing a stalk at the same pole [4]. A stalked cell then elongates its cell body, replicates and segregates its DNA, and produces a flagellum at the pole opposite its stalk prior to cytokinesis. The asymmetric polarization of distinct organelles imparts *C. crescentus* with a highly tractable dimorphic life cycle ideally suited for studying developmental shape changes.

One cell cycle event that requires obvious reshaping of the cell envelope is cell division, which, in nearly all bacteria, requires the conserved tubulin homolog FtsZ. A GTPase, FtsZ polymerizes into a patchy annular structure (the Z-ring) at the incipient division site and recruits the downstream division machinery or divisome. Together, FtsZ and proteins of the divisome coordinate invagination and fission of the membrane(s) with extensive cell wall remodeling [5]. A number of division proteins are known to interact directly with FtsZ. For many of these regulators, however, their mechanism of action toward FtsZ and physiological role in division remain to be discovered. One division complex that has been particularly enigmatic is the ATP-binding cassette (ABC) transporter family complex FtsEX, which is widely conserved in bacteria and functions as a heterodimer with FtsE in the cytoplasm and FtsX in the inner membrane. In *Escherichia coli*, FtsEX localizes to the septum and contributes to the efficiency of cell division, particularly in salt-free media [6]. There is no evidence that FtsEX acts as a transporter, however, and recent studies from a wide range of bacteria have instead implicated FtsEX in the activation of cell wall hydrolysis [7-11]. Septal PG material needs to be split to allow for outer membrane constriction and ultimately for separation of the two new daughter cells. In high salt media, *E. coli* Δ*ftsEX* cells exhibit a phenotype similar to cell wall hydrolysis mutants, i.e. cells are chained and mildly filamentous, suggesting a common genetic pathway [9].

In addition to its involvement in cell wall hydrolysis, in *E. coli*, FtsEX is important for the recruitment of late division proteins and the assembly and/or stability of the septal ring [6]. Interestingly, *E. coli ftsE* mutants impaired for ATP binding and hydrolysis support Z-ring assembly, but constrict poorly [12]. Since FtsE interacts with FtsZ in *E. coli*, one possibility is that FtsEX functions as a membrane anchor for FtsZ and utilizes ATP binding and hydrolysis to regulate Z-ring constriction [13,6].

In this study, we were originally motivated to characterize FtsEX as a novel membrane anchor for FtsZ in *C. crescentus* since FtsE is one of the first proteins recruited to the nascent division site and is important for efficient cell separation and Z-ring assembly and/or stability [14,15]. Considering the conserved function of FtsEX as a modulator of cell wall remodeling, we asked whether FtsEX, in addition to promoting Z-ring structure, regulates cell wall cleavage in *C. crescentus*. We find that *ftsE* has strong synthetic cell separation defects with cell wall hydrolytic factors. Interestingly, however, deleting *ftsE* produces chains of cell bodies connected by thin, tube-like connections that contain all layers of the cell envelope. This is in stark contrast to the thick, uncleaved septa and compartmentalized cytoplasms observed in hydrolysis mutants from *E. coli* and other organisms. The cell-cell connections of Δ*ftsE* cells are, instead, morphologically and topologically similar to *C. crescentus* stalks. In accordance with their shared morphological features, the stalk proteins StpX and PbpC localize to both the skinny connections and stalks of Δ*ftsE* cells, indicating that stalk formation may be mechanistically similar to the elaboration of the extended constriction sites. Our data reveal unanticipated morphogenetic plasticity in *C. crescentus*, with a stalk-like program taking over at failed division sites in the Δ*ftsE* mutant to yield novel cell morphology.

## Results

### *ΔftsE* cells form chains connected by skinny constrictions

To begin to address the role of FtsEX in *C. crescentus*, we first attempted to make *ftsE* and *ftsX* deletion strains. Although *ftsE* is annotated as essential [16,14], we successfully made several independent Δ*ftsE* clones [15]. *ftsX* is also annotated as essential [16], but unlike *ftsE*, we were unable to make an *ftsX* deletion, depletion, or overexpression strain, suggesting that *C. crescentus* cells are highly sensitive to changes in FtsX levels. To understand the role of the FtsEX complex in *C. crescentus* morphogenesis, we focused on characterizing the *ftsE* mutant in detail. Δ*ftsE* cells displayed a striking division phenotype consisting of chained cells with skinny, extended connections between cell bodies. Consistent with the chaining phenotype, Δ*ftsE* cells are longer and grow more slowly than WT [15]. Transmission electron microscopy (TEM) offered us better resolution of cells lacking FtsE (Fig 1). Δ*ftsE* cell bodies were heterogeneous in length, but overall appeared elongated compared to WT, which suggests a delay or inefficiency in the initiation of constriction. The thin connections between Δ*ftsE* cell bodies were also heterogeneous in length, with some extending hundreds of nanometers, and had dimensions qualitatively similar to those observed for stalks (Fig 1B). Overall, the Δ*ftsE* phenotype supports a role for FtsE both in the initiation of constriction and in late stage cell separation.

**Fig 1.**
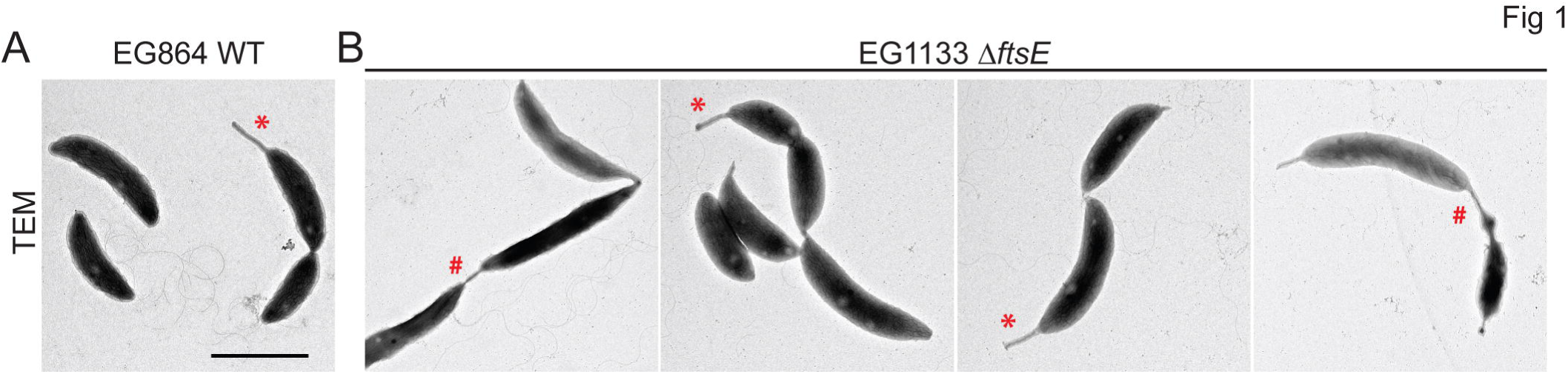
Transmission electron microscopy (TEM) of chained *ΔftsE* cells with thin, extended connections between cell bodies. (A) Micrograph of WT cells. (B) Micrographs of *ΔftsE* cells. * = stalk; # = skinny connection. Scale bar = 2 um.

### FtsE promotes focused Z-ring organization

FtsE has been reported to bind FtsZ in *E. coli*, is one of the first division proteins to localize to midcell after FtsZ in *C. crescentus*, and *C. crescentus* Δ*ftsE* cells have aberrant Z-rings [13-15]. Specifically, in Δ*ftsE* cells FtsZ is more diffuse and often localizes as clusters of puncta instead of focused Z-rings (Fig 2) [15]. These data suggest that FtsE may regulate early Z-ring structure and/or assembly. Consequently, we tested if *ftsE* interacted genetically with the positive Z-ring regulator *zapA*, which, like FtsE, is also recruited early to midcell by FtsZ in *C. crescentus* [17,14]. Δ*zapA*Δ*ftsE* cells displayed mild synthetic growth and cell length defects, but had severely disrupted, diffuse Z-ring structures (Fig 2). We also observed that Δ*zapA*Δ*ftsE* cells were very sensitive to even slight increases in FtsZ levels and were noticeably filamentous after only one hour of *ftsZ-cfp* expression (Fig 2D). We conclude that FtsE contributes to proper Z-ring focusing at midcell and that the elongated cell bodies in Δ*ftsE* are likely due to inefficient initiation of constriction by the aberrant FtsZ structures.

**Fig 2.**
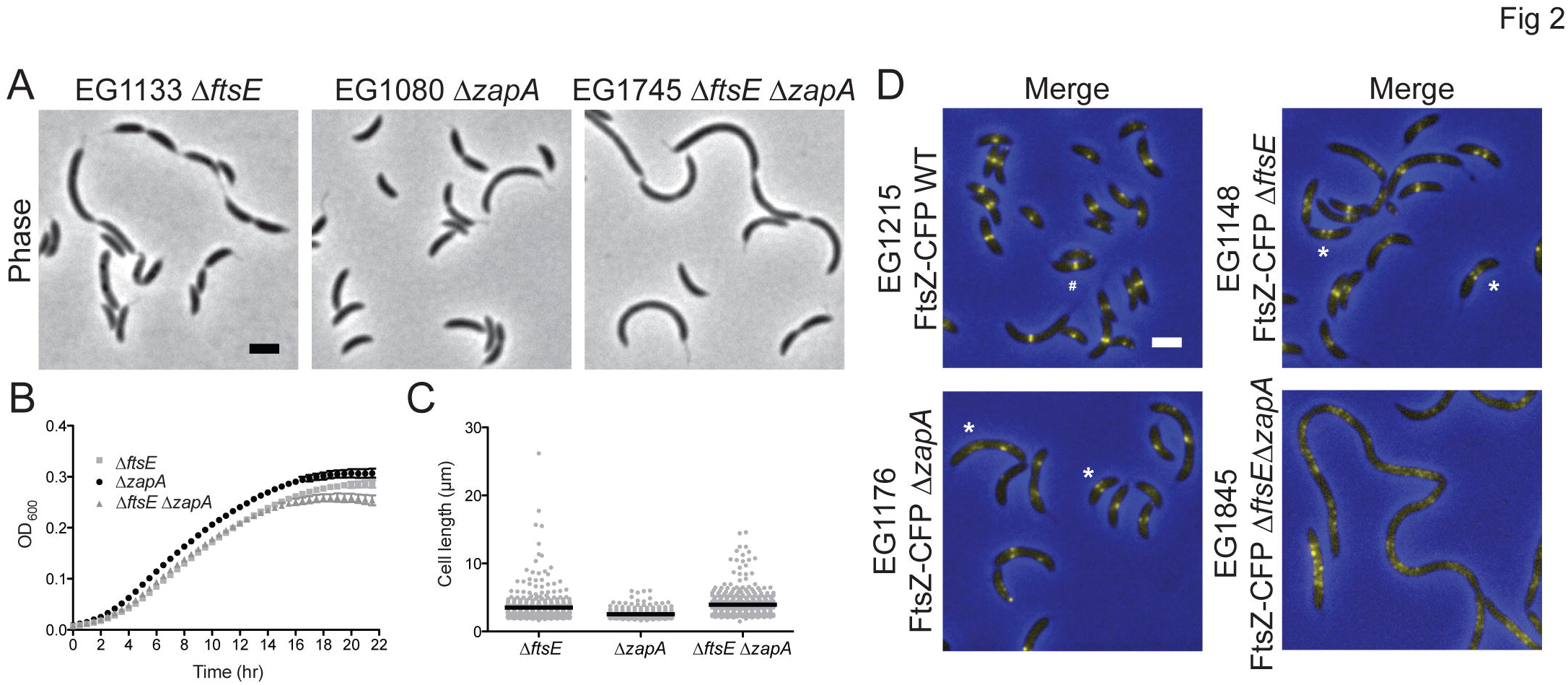
*ftsE* has synthetic growth, cell length, and Z-ring structural defects with Z-ring regulator *zapA*. (A) Phase contrast micrographs of Δ*ftsE*,Δ*zapA*, and Δ*ftsE*Δ*zapA* cells. (B) Growth curves of strains in (A). Doubling time (h) ± SEM: EG1080 = 1.78 ± 0.01; EG1133 = 1.81 ± 0.02; EG1745 = 1.86 ± 0.02. Differences between strain doubling times were ns by one-way ANOVA. (C) Cell length analyses of strains in (A). Mean cell length (μm) ± SEM: EG1080 = 2.54 ± 0.03; EG1133 = 3.50 ± 0.11; EG1745 = 3.94 ± 0.10. All pairwise comparisons of mean cell lengths yielded p values < 0.01 by one-way ANOVA. (D) FtsZ-CFP localization after 1 h of induction in WT, ΔftsE,Δ*zapA*, and Δ*ftsE*Δ*zapA* cells. # = focused Z-ring; * = diffuse, punctate Z-ring. Scale bars = 2 μm.

### Excess FtsE or FtsEX alters the localization of FtsZ and new cell wall synthesis

Because cells lacking FtsE have perturbed Z-ring organization, we hypothesized that overproducing FtsE would affect Z-ring structure, particularly if FtsE binds directly to FtsZ. Overexpression of either *ftsE* or *ftsEX* caused dramatic filamentation, and overexpression of *ftsE* alone also caused ectopic poles to form (Fig 3). After four hours of FtsE overproduction, instead of Z-rings, FtsZ-CFP formed discrete puncta along the length of the filamentous cells (Fig 3A). Interestingly, when we overproduced FtsEX, FtsZ-CFP localized in a drastically different pattern, as multiple wide bands (Fig 3A). *C. crescentus* Z-ring positioning is in part dictated by a negative regulator of FtsZ assembly called MipZ, which forms a complex near the origin of replication [18]. After the polar origin region is duplicated, the second copy is quickly transported to the opposite cell pole. Bipolar MipZ thereby directs Z-ring assembly at midcell by inhibiting FtsZ polymerization at the poles [18]. MipZ-YFP localized at the poles and as fairly regularly spaced puncta in cells overproducing FtsE or FtsEX, but its localization was more diffuse in cells overproducing FtsEX (Fig 3B). We interpret the MipZ localization data as evidence that chromosomal replication and segregation still occur in cells overproducing FtsE despite the inhibition of division; however high levels of FtsEX may interfere with levels or localization of MipZ.

**Fig 3.**
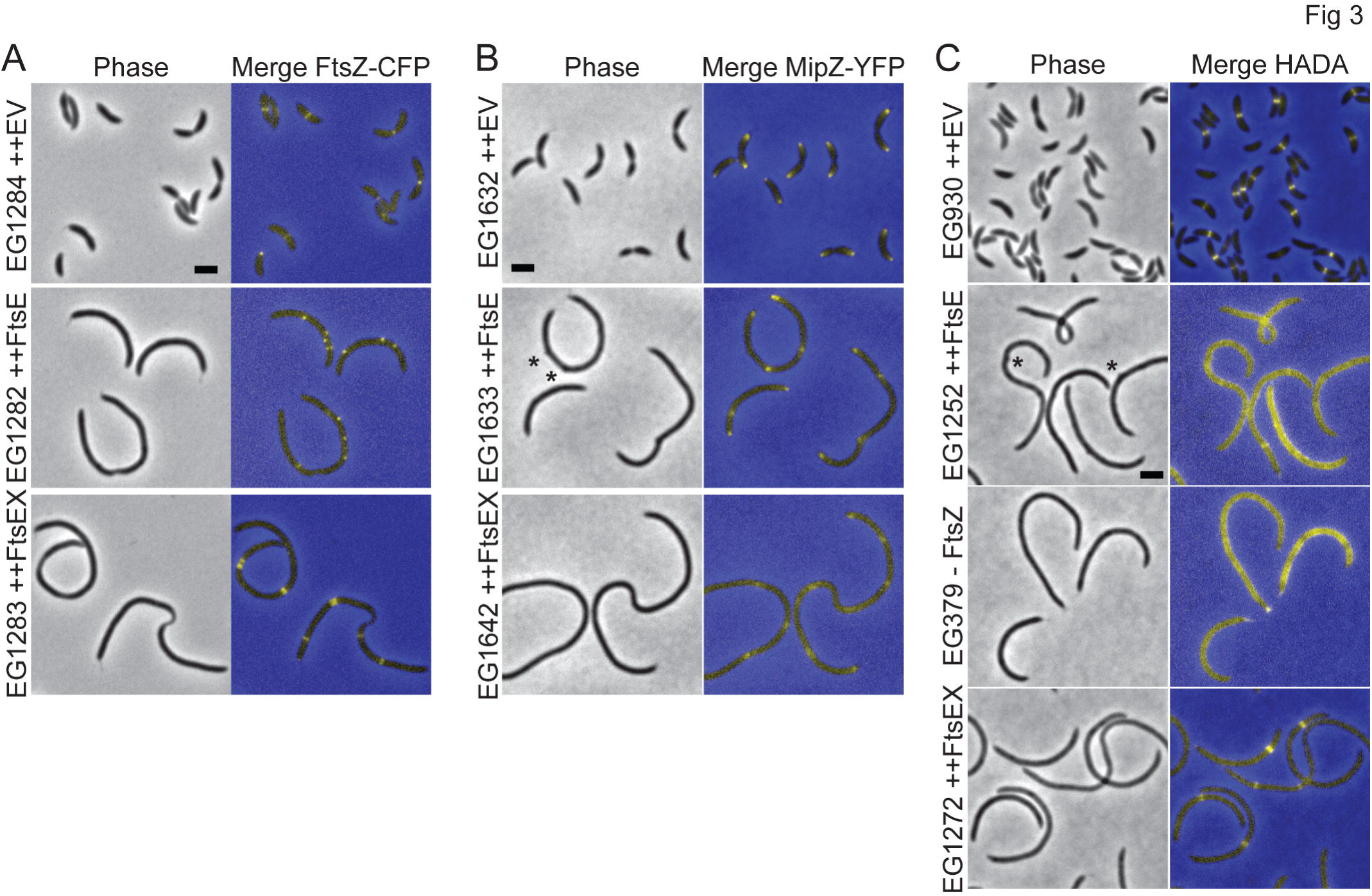
Overproducing FtsE or FtsEX causes filamentation and affects the localization of FtsZ and new PG. (A) FtsZ-CFP localization after 1 h of induction in cells bearing an empty vector (EV) or overexpressing *ftsE* or *ftsEX* for 4 h. (B) MipZ-YFP localization in cells bearing and EV or overexpressing *ftsE* or *ftsEX* for 4 h. (C) HADA labeling of cells depleted for FtsZ for 4 h, bearing an EV, or overexpressing *ftsE* or *ftsEX* for 4 h. * = ectopic poles. Scale bars = 2 μm.

In *C. crescentus*, FtsZ directs new cell wall synthesis at midcell even before the onset of division (Fig 3C) [19]. To determine if the FtsZ structures in FtsE or FtsEX overproducing cells were competent to localize new cell wall synthesis, we pulse-labeled cells with the fluorescent D-amino acid hydroxycoumarin-carbonyl-amino-D-alanine (HADA), which can replace D-Ala^4^ or D-Ala/Gly^5^ in the lipid II peptide side chain, to track the incorporation of newly synthesized PG [20]. High levels of FtsE resulted in diffuse HADA labeling which closely resembled the pattern of HADA incorporation in FtsZ-depleted cells (Fig 3C). This result suggests that the FtsZ puncta associated with *ftsE* overexpression are unable to direct local cell wall metabolism. Cells overexpressing *ftsEX*, however, had discrete, wide bands of new PG incorporation, likely directed by the similarly organized Z-ring structures (Fig 3C). Thus, high levels of FtsE or FtsEX not only differentially affect Z-ring organization, but also affect FtsZ’s ability to locally incorporate new cell wall material at midcell. Specifically, the intact FtsEX complex is required both for formation of Z-rings and downstream communication with PG synthetic machinery.

### *ftsE* interacts genetically with PG hydrolytic factors and a regulator of stalked pole development

Since FtsEX has been implicated in cell wall hydrolysis in numerous bacteria and Δ*ftsE* cells displayed a cell chaining phenotype, we asked whether FtsEX, in addition to promoting Z-ring structure, regulates *C. crescentus* cell wall hydrolysis. In *E. coli*, periplasmic *N*-acetylmuramyl-_L_-alanine amidases AmiA/B/C are responsible for cleaving bonds that link stem peptides to glycan strands at the septum [21]. The amidases require activation by the LytM domain containing proteins EnvC, which stimulates AmiA/B, and NlpD, which stimulates AmiC, to split apart septal PG [22,23]. FtsEX directly recruits EnvC to the septum via the periplasmic extracellular loop (ECL) of FtsX and the coiled coil (CC) domain of EnvC [9]. *C. crescentus* possesses a limited number of lytic enzymes involved in peptidoglycan remodeling, with only a single *N*-acetylmuramyl-_L_-alanine amidase, most similar to *E. coli* AmiC [24,25]. There are at least seven genes coding for putative LytM domain containing proteins; however, the only characterized protein in the *C. crescentus* LytM family is DipM, which participates in cell wall remodeling and coordinated constriction of the cell envelope layers at the division plane [25-27]. To determine whether the FtsEX PG hydrolysis paradigm applies to *C. crescentus*, we performed a BLAST search for *E. coli* EnvC homologues and found CCNA_03547, which we will hereafter refer to as *L*ytM domain protein *F* (LdpF) (Martin Thanbichler, personal communication). Like EnvC, LdpF is a LytM domain containing protein with a signal peptide, two N-terminal CC domains, and a C-terminal LytM domain (Fig S1A). We hypothesized that *C. crescentus* FtsEX-LdpF-AmiC may function in an activation pathway analogous to *E. coli* FtsEX-EnvC-AmiA/B.

We first adopted a genetic approach to investigate the role of FtsEX in cell wall hydrolysis during division. In *E. coli*, FtsEX is required for EnvC’s localization at midcell [9]. However, LdpF-mCherry is diffuse, and we did not observe differences in its localization between WT and Δ*ftsE* cells (Fig S1B). Although *E. coli* cells lacking EnvC or AmiA/B are not as sick as cells lacking FtsEX, part of the division defect associated with loss of FtsEX may be due to EnvC inactivation [9]. Consistent with this reasoning, in *C. crescentus*, cells lacking LdpF or AmiC had mild chaining and growth defects compared to Δ*ftsE* cells (Fig 4). Depleting AmiC in Δ*ftsE* or Δ*ldpF* backgrounds caused strong synthetic cell separation and growth defects, accompanied by noticeable lengthening of the skinny connections between cell bodies, particularly in the *ftsE* mutant (Fig 4). Combining Δ*ldpF* with loss of FtsE produced only mild synthetic growth and chaining defects, which is consistent with them acting in a common pathway (Fig 4).

**Fig 4.**
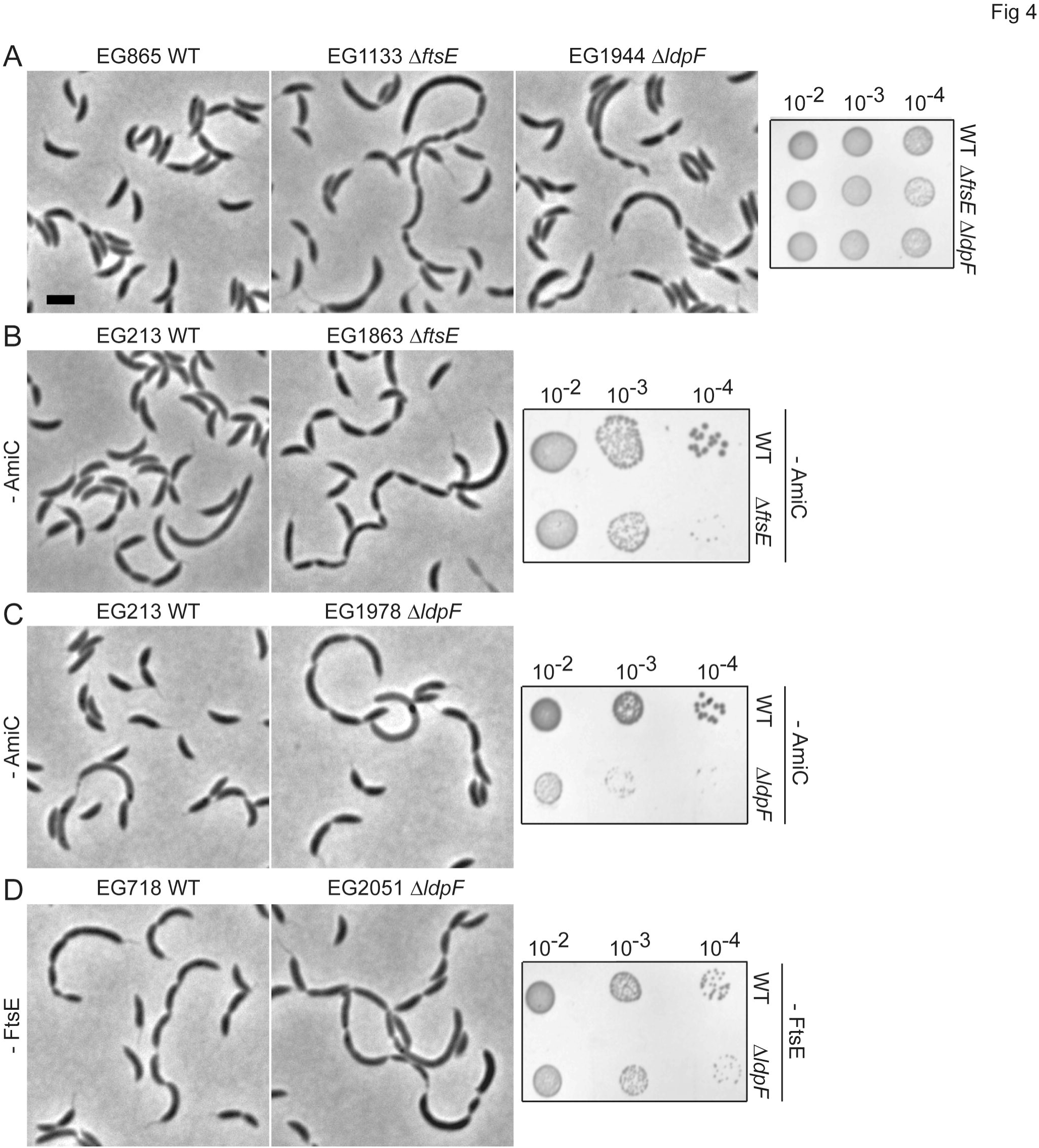
Synthetic genetic interactions between *amiC, ftsE*, and *ldpF*. Phase contrast micrographs and serial spot dilutions of (A) WT, Δ*ftsE*, and Δ*ldpF*; (B) WT and Δ*ftsE* cells depleted for AmiC for 24 h; (C) WT and Δ*ldpF* cells depleted for AmiC for 24 h; (D) WT and Δ*ldpF* cells depleted for FtsE for 24 h. Scale bar = 2 μm.

*E. coli* cells lacking EnvC depend on NlpD for cell separation: simultaneous inactivation of either EnvC or FtsEX and NlpD results in severe chaining, supporting the hypothesis that FtsEX activates EnvC’s ability to promote septal PG cleavage [22]. In *C. crescentus*, the LytM protein most closely related to NlpD is DipM, which has similar domain organization to NlpD but differs in that it is not associated with the outer membrane. Depleting AmiC in cells lacking DipM caused a moderate synthetic growth defect, whereas depletion of DipM in Δ*ftsE* or Δ*ldpF* cells was synthetic lethal (Fig 5). The synthetic defects associated with loss of DipM with FtsE, LdpF, or AmiC as well as the distinct morphology of Δ*dipM* cells [25-27] suggests that DipM operates in a non-redundant, parallel hydrolytic pathway. We also hypothesize, based on the strong synthetic interactions between AmiC and FtsE or LdpF but only mild synthetic defects associated with loss of FtsE and LdpF, that AmiC operates in a hydrolytic pathway separate from FtsE and LdpF. However, it is apparent that all three putative PG hydrolytic pathways contribute to the efficiency of cell separation in *C. crescentus*.

**Fig 5.**
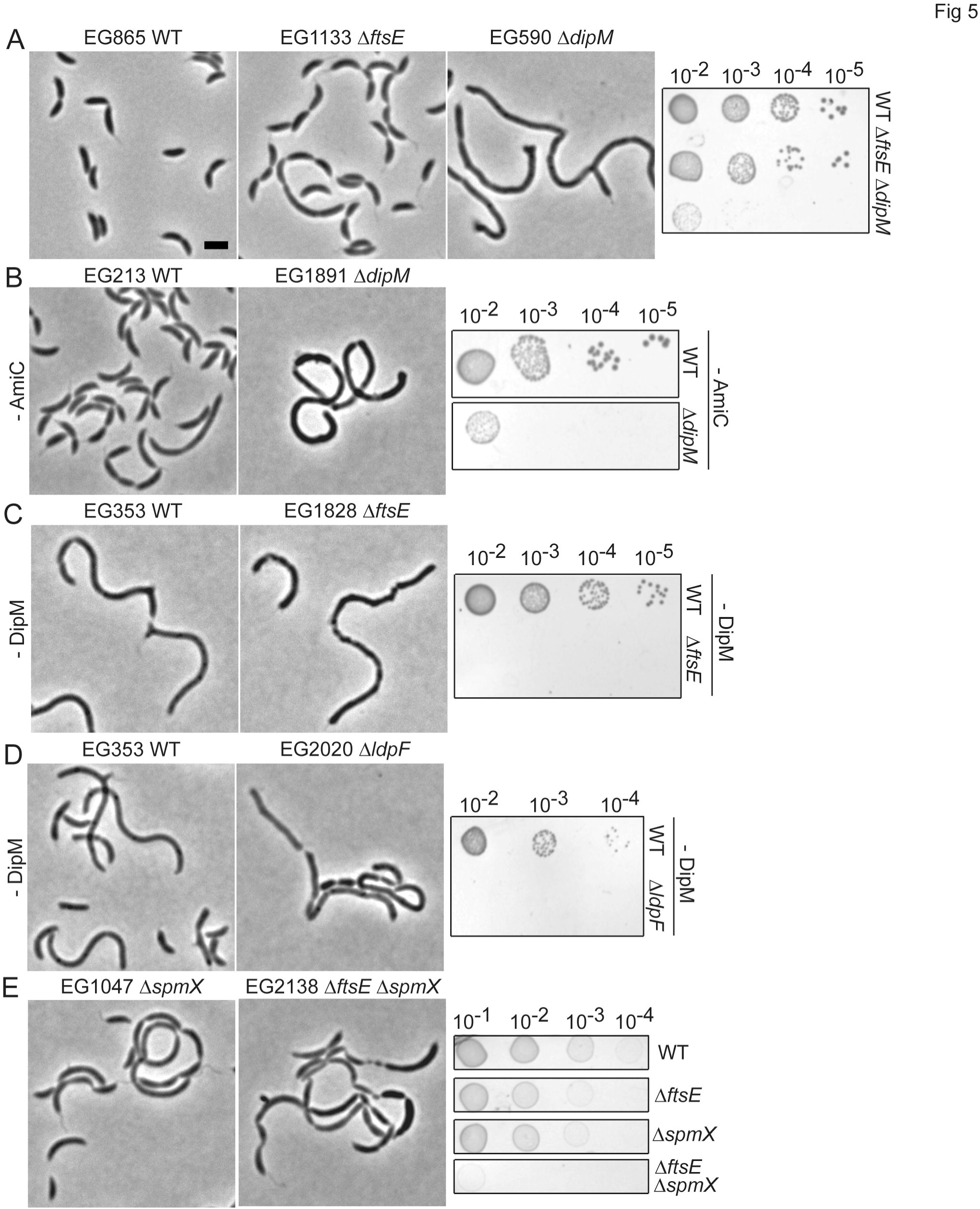
Synthetic genetic interactions between *dipM* and *amiC, ftsE*, or *ldpF* and between *ftsE* and *spmX*. Phase contrast micrographs and serial spot dilutions of (A) WT, Δ*ftsE*, and Δ*dipM;* (B) WT and Δ*dipM* cells depleted for AmiC for 24 h; (C) WT and Δ*ftsE* cells depleted for DipM for 24 h; (D) WT and Δ*ldpF* cells depleted for DipM for 24 h; (E) Δ*spmX* and Δ*ftsE*Δ*spmX*. Scale bar = 2 μm.

Interestingly, in a transposon deep-sequencing analysis to uncover transposon insertions that alter the competitive fitness of Δ*spmX* cells, the *ftsE* and *ldpF* loci were the first and second least favored insertion sites, respectively, in Δ*spmX* cells as compared to WT [28]. We therefore tested for synthetic interactions between *ftsE* and *spmX*, which encodes a muramidase homolog that localizes to the stalked cell pole and controls the swarmer-to-stalk cell transition [29]. Consistent with the Δ*spmX* transposon deep-sequencing results, we observed severe synthetic growth and morphological defects for Δ*ftsE*Δ*spmX* cells, implicating FtsE and LdpF in the stalked pole development pathway, likely through a connection to cell wall remodeling (Fig 5E).

### The CC domain of LdpF interacts with the ECL of FtsX, but LdpF does not activate AmiC PG hydrolysis *in vitro*

To provide biochemical support for our *in vivo* findings, we purified LdpF, AmiC, DipM, and the ECL of FtsX, and monitored PG degradation using an *in vitro* dye release assay [30,23]. Although bacterial two hybrid showed a positive interaction for the ECL of FtsX and the CC domain of LdpF, we did not observe LdpF-activated AmiC PG hydrolysis *in vitro* (Fig S2; S1 Text). Interestingly, the LytM domain of DipM was sufficient to mildly stimulate AmiC hydrolase activity *in vitro* (Fig S2C; S1 Text). However, as *C. crescentus* DipM and AmiC are most similar to the *E. coli* activator-amidase pair, NlpD and AmiC, perhaps this result is unsurprising. In light of the distinct phenotypes of cells lacking DipM and AmiC, however, we suspect that DipM has activities in addition to the regulation of AmiC. Collectively, our genetic and biochemical evidence indicate at least three hydrolytic pathways in *C. crescentus* and implicate a yet unidentified downstream target of FtsEX-LdpF in the regulation of PG metabolism.

### Electron cryotomography reveals unique cell envelope organization at chaining sites

Our genetic evidence is consistent with a role for FtsEX in regulating cell wall hydrolysis for cell separation. Although TEM highlighted the general cell separation defects of Δ*ftsE* cells (Fig 1), electron cryotomography (ECT) allowed us to dissect the exact stages at which these cells are blocked during division (Fig 6). To capture the cell envelope organization at the skinny cell-cell connections, we imaged cells lacking both FtsE and AmiC since loss of AmiC exacerbates the Δ*ftsE* chaining phenotype (Fig 4). In five out of six tomograms, we could identify with certainty the presence of all layers of the cell envelope in the skinny connections between chained cell bodies (Fig 6). Four of these had obvious cytoplasmic volume between the unfused inner membranes. In at least one example, however, the inner membranes were closely stacked on top of each other, but not fused (Fig 6).

**Fig 6.**
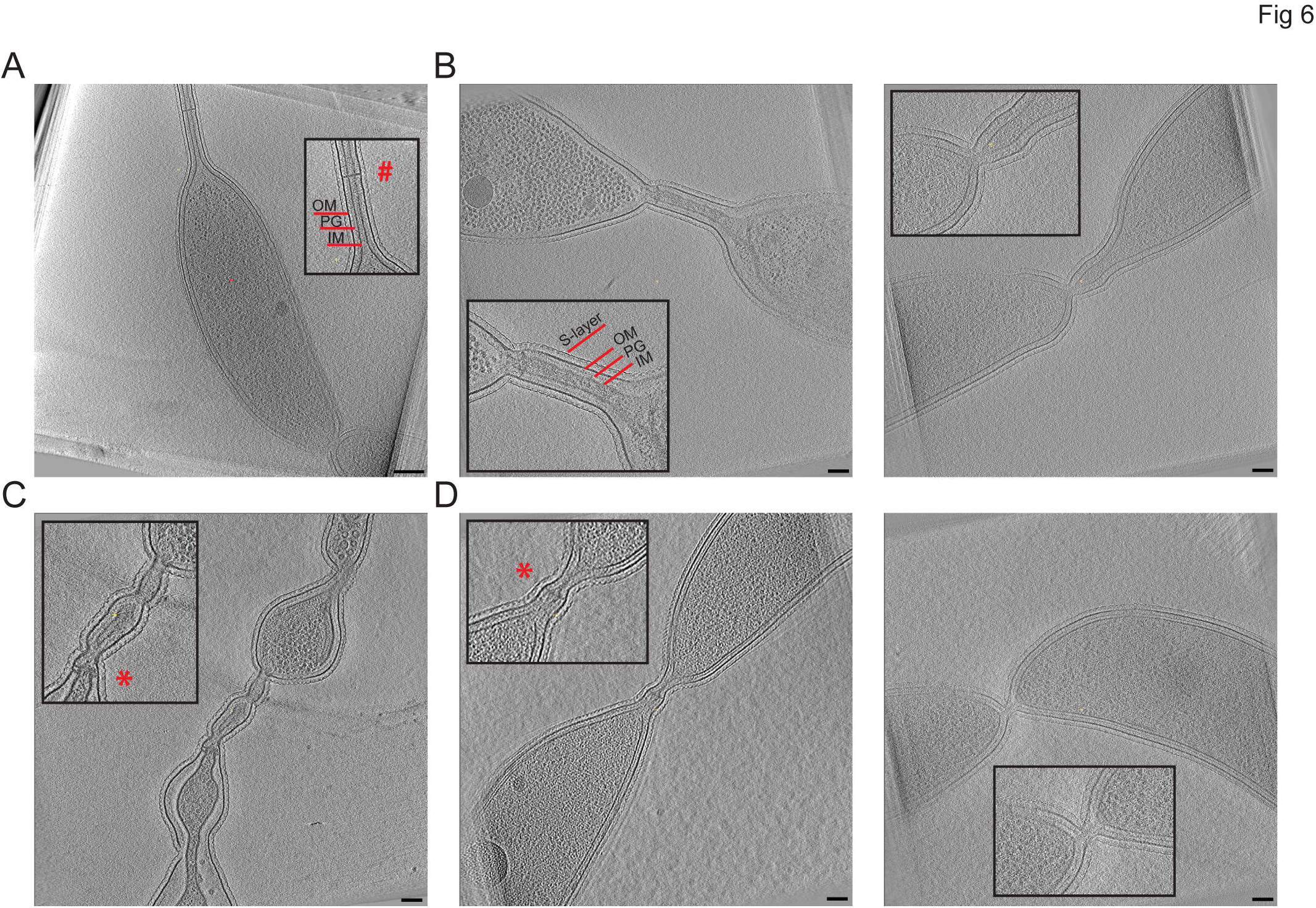
Electron cryotomography (ECT) of cells lacking FtsE and AmiC reveal stalk-like connections between cell bodies. (A) WT *C. crescentus* stalk with cross-bands. (B) *ftsE* mutants with skinny connections most similar to stalks. (C) *ftsE* mutants with skinny connections that are stalk-like, but that have regions with heterogeneous widths. (D) *ftsE* mutants with skinny connections that are fairly short with inner membranes that are close together or nearly fused. Abbreviations are as follows: OM = outer membrane, PG = peptidoglycan, IM = inner membrane. *#* = cross-band; * = cross-band-like structure. Scale bar (A) = 200 nm; Scale bars (B-D) = 100 nm.

In WT *C. crescentus*, the final stages of inner membrane fission are rapid, and the smallest diameter for inner membrane connections that have been captured by ECT are ∼60 nm [31]. In cells lacking FtsE and AmiC with fairly uniform connections (Fig 6B, D), we observed inner membrane diameters ranging from ∼12 to 60 nm. Others were more variable and had intermittent bulging, with inner membrane diameters ranging from ∼20 to 300 nm within a single cell-cell connection (Figs 6C, S3). This organization of the cell envelope is strikingly different from other mutants deficient in cell wall hydrolysis.

*E. coli* cells lacking EnvC and NlpD or all four LytM-domain containing factors complete inner membrane fusion and cytoplasmic compartmentalization, but struggle to constrict their outer membrane due to a layer of intact PG between adjacent chained cells [22]. Similarly, Δ*dipM* cells form chains with fused inner membranes and thick, multilayered PG between cell bodies [25,26]. Therefore, the cell envelope organization of cells lacking FtsE and AmiC, namely the continuous cytoplasmic connections and narrow spacing between the inner membranes, represents a unique cell separation phenotype and, potentially, a novel pathway for cell separation in *C. crescentus*.

### Δ*ftsE* thin connections are morphologically stalk-like and enriched for stalk proteins

During its dimorphic life cycle, *C. crescentus* elaborates a polar stalk, a tubular extension of the cell envelope important for nutrient uptake [32]. ECT of cells lacking FtsE and AmiC reinforced an observation we had previously made based on TEM of Δ*ftsE*, namely, the striking morphological similarities between the extended connections of *ftsE* mutants and *C. crescentus* stalks (Figs 1, 6). In addition to sharing approximate widths (∼12-300 nm inner membrane diameter for the skinny connections; ∼20-40 nm inner membrane diameter for the stalks) and cell envelope organization, we occasionally observed electron dense structures that spanned the short axis of the cell envelope of the thin connections (Fig 6). These structures were reminiscent of stalk cross-bands, multiprotein assemblies that transect the stalk at regular intervals and function as diffusion barriers to compartmentalize stalk and cell body periplasmic and membrane proteins (Figs 6A,C, D, S3) [33].

Since stalk growth occurs by incorporation of new material at the cell body-stalk junction [32], we monitored HADA incorporation at the extended constrictions of *ftsE* mutants. In general, we observed incorporation of new cell wall material throughout the skinny connections (Fig S4), which contrasts with the pattern of *de novo* PG synthesis only at the base of stalks. There are two modes of zonal PG synthesis in *C. crescentus*: FtsZ-independent PG incorporation at the base of the stalk and FtsZ-dependent PG incorporation at midcell [19]. Interestingly, in Δ*ftsE* cells, FtsZ is not enriched at the skinny connections suggesting that the new cell wall synthesis occurring throughout the skinny connections is an FtsZ-independent process similar to PG synthesis at the base of the stalk (Fig S4).

Motivated by the morphological similarities between the extended constrictions in Δ*ftsE* cells and *C. crescentus* stalks, we asked if any stalk-specific proteins could localize to the skinny connections. StpX is a bitopic membrane protein enriched in the stalk that regulates stalk length [34]. We expressed *stpX-cfp* in WT, Δ*ftsE*, and Δ*ftsE* cells lacking AmiC: StpX-CFP localized to the stalks in all genetic backgrounds, but strikingly, was also enriched at the skinny constrictions in the *ftsE* mutant cells (Fig 7A). The bifunctional penicillin binding protein, PbpC, is involved in stalk elongation [35] and is required to sequester StpX at the stalk in WT *C. crescentus* [37]. To investigate the localization dependency of StpX to the skinny constrictions, we expressed *stpX-cfp* in a Δ*ftsE*Δ*pbpC* mutant background. Consistent with previous reports, StpX-CFP was not enriched in stalks in cells lacking both FtsE and PbpC. However, StpX-CFP was also not enriched at the skinny connections between cells and instead was diffusely localized at the cell periphery (Fig 7B). We also expressed *pbpC-yfp* in WT and Δ*ftsE* cells and observed enrichment at the base of stalks as well as frequent localization to the skinny connections in the *ftsE* mutant (Fig 7C). Since we observe enrichment of both PbpC and StpX at the skinny connections in Δ*ftsE* cells, we hypothesize that PbpC initially recruits StpX to the skinny connections and promotes its retention at those sites.

**Fig 7.**
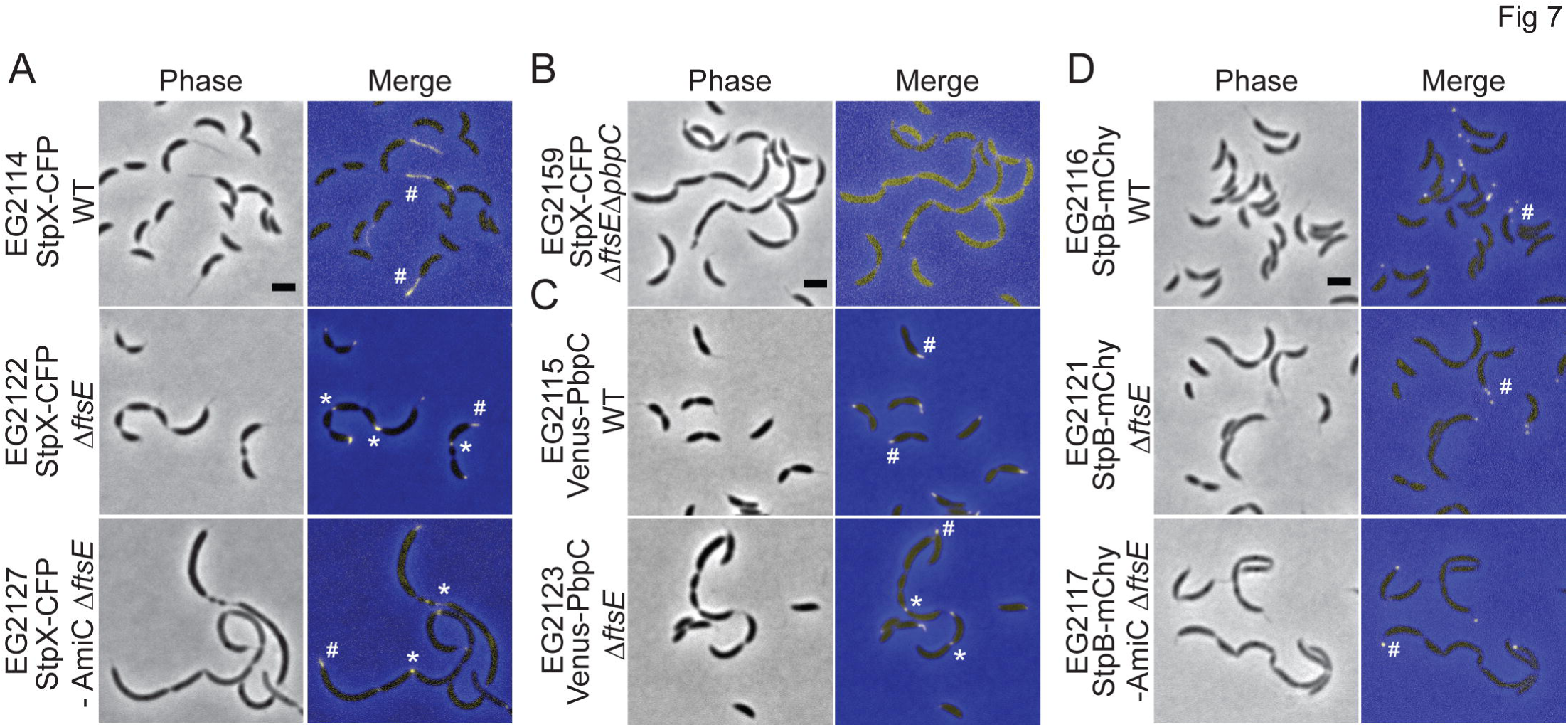
Stalk specific proteins StpX and PbpC localize to the skinny constrictions in *ftsE* mutants. (A) Localization of StpX-CFP induced for 18 h in WT, Δ*ftsE* or Δ*ftsE* cells depleted for AmiC for 18 h. (B) Localization of StpX-CFP induced for 2.75 h in Δ*ftsE*Δ*pbpC*. (C) Localization of Venus-PbpC induced for 2 h in WT or Δ*ftsE*. (D) Localization of cross-band protein StpB-mCherry induced for 18 h in WT, Δ*ftsE* or Δ*ftsE* cells depleted for AmiC for 18 h. # = stalk enrichment; * = skinny connection enrichment. Scale bars = 2 μm.

Considering the presence of both PbpC and StpX at the skinny connections in the *ftsE* mutant, we monitored the localization of the stalk cross-band protein, StpB. StpB-mCherry localized as puncta in the stalks of WT, Δ*ftsE*, and Δ*ftsE* cells lacking AmiC, but was not enriched at the extended constriction sites (Fig. 7D). Consequently, the proteinaceous, envelope-spanning discs observed in the *ftsE* mutant by ECT (Fig 6) may not, in fact, be cross-bands or may differ in molecular composition from stalk cross-bands. We conclude that the skinny connections share numerous morphological and molecular similarities with stalks, but the two structures are not physically or biochemically identical.

## Discussion

The role of FtsEX in synchronizing PG remodeling with cell division appears to be conserved amongst distantly related bacterial species such as *E. coli, S. pneumoniae*, and *M. tuberculosis*, although the downstream adaptor or enzyme targets vary [7-11]. We provide evidence that this paradigm also extends to the α-proteobacterium *C. crescentus*. Our data indicate that FtsE is important for initial Z-ring assembly and regulates Z-ring structure in a manner dependent on its stoichiometry with FtsX (Figs 2, 3, 8A). Different levels of FtsE or FtsEX not only affect FtsZ localization, but also FtsZ function, namely its ability to localize incorporation of new cell wall material (Fig 3). Additionally, our data implicate FtsEX in a cell wall metabolic pathway involving LdpF and an unidentified downstream cell wall factor regulated by LdpF (Fig 4, 8B). Thus *C. crescentus* FtsEX, similar to what has been proposed in *E. coli*, may synchronize PG remodeling with Z-ring constriction during division [9]. ECT of the skinny connections in Δ*ftsE* revealed a cell envelope architecture remarkably distinct from *E. coli* hydrolase mutants, however, and an overall morphology that was strikingly stalk-like (Fig 6). Enrichment for stalk proteins StpX and PbpC at the skinny constrictions in Δ*ftsE* and a strong genetic interaction between *ftsE* and the stalked cell fate determinant *spmX* further support mechanistic overlap between the elaboration of thin connections in *ftsE* mutants and stalk development (Figs 5, 7, 8A).

**Fig 8.**
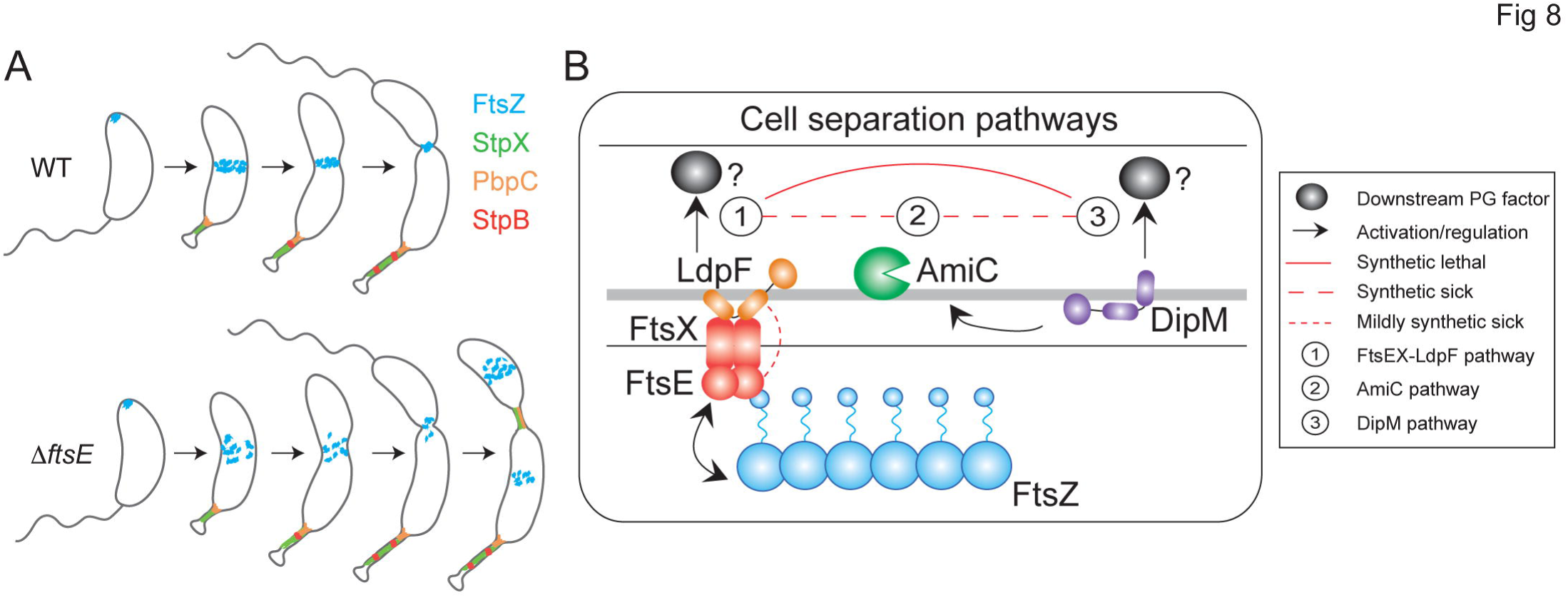
FtsEX-mediated regulation of constriction initiation and final cell separation reveals morphogenetic plasticity in *C. crescentus*. (A) Cell cycle localization of FtsZ and stalk proteins in WT and Δ*ftsE* cells. (B) Three putative cell separation pathways in *C. crescentus*.

During the late stages of division in *C. crescentus*, constriction of the inner membrane proceeds until the inner membranes of the two future daughter cell compartments are connected only by a small tubular structure [31]. Out of thousands of cells Judd and colleagues examined in their study, only five displayed inner membranes with diameters less than ∼100 nm and the smallest inner membrane connection was 60 nm in diameter [31]. ECT of cells lacking FtsE and AmiC with fairly uniform connections showed inner membrane diameters ranging from ∼12 to 60 nm (Fig 6). Thus, the majority of cells lacking FtsE and AmiC have inner membrane diameters that fall well below the lowest threshold reported for inner membrane diameters at any stage of WT cell division. Furthermore, WT cells spend a short amount of time in these late, transitional stages, perhaps only a few seconds, and membrane topology changes very rapidly [31]. In the case of *ftsE* mutant cells, the thin connections are abundant in a mixed cell population and, based on their nearly abutting inner membranes, are likely blocked or delayed at a terminal stage just prior to inner membrane fission. In WT *C. crescentus*, the mechanisms underlying rapid terminal constriction and membrane fusion are unknown [31]. Our data indicate that FtsEX is important for inner membrane fusion, either directly or indirectly, and that in the absence of FtsE, inner membrane fusion frequently fails, PG synthesis continues, and cells elaborate a stalk-like structure.

Understanding how proteins are targeted specifically to the stalk is important for understanding mechanisms of subcellular organization in bacteria as well as stalk function [36]. We have limited knowledge about the molecular pathways responsible for targeting proteins to the stalk; however, stalked pole geometry, membrane curvature, or unique peptidoglycan motifs are possible mechanisms for protein localization at the stalk. The observation that both StpX and PbpC localize to the skinny connections in Δ*ftsE* cells provides an experimental handle for understanding stalk biogenesis and stalk protein localization cues. In WT *C. crescentus*, the bactofilins, BacA and BacB, localize at the stalked pole and recruit PbpC during the swarmer-to-stalked cell transition [35]. We have not monitored the localization of the bactofilins in Δ*ftsE* cells, but we predict that they would localize at the skinny connections upstream of PbpC, similar to the protein recruitment hierarchy observed for stalks. It has been proposed that the membrane curvature at the stalk-cell body junction drives BacAB clusters to localize there [35]. These BacAB clusters could likewise recruit PbpC and StpX to the junctions between the cell bodies and skinny connections in Δ*ftsE* on the basis of shared membrane curvature with the cell body-stalk junction. Additionally, the composition of the peptidoglycan is purportedly distinct between the cell body and the stalk [37], as stalks are more resistant to lysozyme treatment [38]. PbpC may regulate the rigidity of the stalk by dictating a specific cell wall remodeling regime, which may also be active at the skinny connections [35]. The periplasmic N-terminal domain of StpX is required for its stalk localization and may recognize PbpC-specific changes in PG chemistry, leading to sequestration of StpX in the stalk and the skinny constrictions of Δ*ftsE* cells [34].

Within α-proteobacteria, asymmetric patterns of growth are particularly well-represented in the orders *Rhizobiales* and *Caulobacterales* [39]. Unlike *C. crescentus*, which divides by asymmetric binary fission, *Rhodomicrobium vannelliei* and *Hyphomonas neptunium*, members of *Rhizobiales* and *Caulobacterales* respectively, use a budding mechanism whereby new offspring emerges from the tip of a stalk structure [40,39]. Cell division thus occurs in an extremely asymmetric manner at the bud neck, producing a stalked mother cell and a non-stalked daughter cell [39]. After initially increasing in cell size by dispersed PG incorporation, *H. neptunium* displays a period of zonal growth at the new cell pole leading to the elaboration of a stalk structure [39]. This type of stalk outgrowth is similar to what occurs in other stalked α-proteobacteria like *C. crescentus*, which suggests conservation of core machinery. The asymmetric manner in which *H. neptunium* divides exemplifies how stalks may function not only as specialized organelles, but also as division planes, depending on the bacterial species [39]. The morphology of Δ*ftsE* cells, with thin, stalk-like extensions between cell bodies is reminiscent of a predivisional *H. neptunium* cell where the stalk is closely integrated within the cell division program.

While specialized protein machineries exist for cell division and cell elongation, there is no single protein, much less entire machinery, absolutely and specifically required for stalk formation. Since the elongation machinery proteins MreB and RodA contribute to stalk growth and morphogenesis, stalk synthesis has been proposed to be a specialized form of cell elongation [41]. Thus, it is possible that the skinny, stalk-like connections between cell bodies in Δ*ftsE* are a result of a modified form of cell elongation, similar to what has been postulated for stalks [41]. Our data suggest that although the slender connections in Δ*ftsE* share certain morphological and molecular features with stalks, they are not, in actuality, ectopic stalks forming at failed division sites: the spatial pattern of PG incorporation is distinct from stalks, the diameters are not as homogeneous as for stalks, and StpB does not localize at the skinny connections (Figs S3, 6, 7). We instead favor the hypothesis that when Δ*ftsE* cells stall at a late stage in division, the division machinery disengages and the cell elongation machinery takes over and elaborates a thin, stalk-like connection. We interpret this phenomenon as an example of morphogenetic plasticity, whereby small changes to established morphogenetic machineries give rise to novel or, in the case of Δ*ftsE*, modified forms of preexisting structures. Overall, our findings have important implications for understanding late stage division regulation, stalk formation, and the coordination of morphogenetic events and machineries in *C. crescentus*. Loss of FtsE has revealed unexpected morphogenetic plasticity between the division and stalk synthesis programs and offers insight into the geneses of diverse morphologies in bacteria.

## Materials and Methods

### Growth conditions for bacterial strains

*C. crescentus* NA1000 strains were grown in peptone yeast extract (PYE) medium at 30^°^C [42]. Additives and antibiotics were used at the following concentrations in liquid (solid) media for *C. crescentus:* xylose 0.3 (0.3)%, glucose 0.2 (0.2)%, vanillate 0.5 (0.5) mM, gentamycin 1 (5) μg ml^-1^, kanamycin 5 (25) μg ml^-1^, spectinomycin 25 (100) μg ml^- 1^, streptomycin (5 μg ml^-1^). Before changes in induction conditions, cells were washed two to three times in plain media. Growth rate analyses were performed in 96-well plates with shaking at 30^°^C using a Tecan Infinite 200 Pro plate reader. Strains and plasmids used in this study are included as Supporting Information Table S1.

### Light microscopy and image analysis

Cells were imaged during the log phase of growth after immobilization on 1% agarose pads. Light microscopy was performed on a Nikon Eclipse Ti inverted microscope equipped with a Nikon Plan Fluor x 100 (numeric aperture 1.30) oil Ph3 objective and Photometrics CoolSNAP HQ cooled CCD (charge-coupled device) camera. Chroma filter cubes were used as follows: ET-EYFP for YFP and ET-ECFP for CFP, ET-dsRED for mCherry and ET-ECFP for HADA. Images were processed in Adobe Photoshop. Automated cell length analysis was performed using Oufti or MicrobeTracker [43]. In MicrobeTracker, algorithm 4 was used for determining cell outlines, with the following parameter change: areaMin=150.

### Whole cell TEM

Cells from EG864 and EG1133 were grown in PYE and prepared for whole cell TEM exactly as described [42].

### HADA labeling

Cells from strains EG213, EG379, EG864, EG930, EG1133, EG1252, EG1272, and EG1863 were grown in PYE. HADA [20] was added to 0.41 mM and the cultures were returned to the shaker for 5 min. The cells were then washed twice with PBS and resuspended in PBS before imaging [42].

### Bacterial two-hybrid

The T18 and T25 plasmids were co-transformed into BTH101 (*F-, cya-99, araD139, galE15, galK16, rpsL1* (*Str ^r^*), *hsdR2, mcrA1, mcrB1*; Euromedex) competent cells, plated onto LB agar with ampicillin (100 μg/μl) and kanamycin (50 μg/μl), and incubated overnight at 30^°^C. Several colonies were inoculated into LB with ampicillin (100 μg/μl), kanamycin (50 μg/μl), and IPTG (0.5 mM) and incubated at 30^°^C overnight. The next morning 2 μl of each culture was spotted onto plates containing ampicillin, kanamycin, X-gal (40 μg/ml), and IPTG (0.5 mM) and incubated for 1-2 days at 30^°^C. Positive interactions were indicated by blue colonies. Every interaction was tested in triplicate.

### Electron cryotomography (ECT)

For ECT imaging, strain EG1863 was grown in PYE with xylose. Once in log phase, EG1863 was washed twice with PYE and resuspended in PYE without xylose to deplete AmiC. EG1863 was then grown in PYE without xylose for ∼6 h, transferred to an eppendorf tube, and shipped on ice to Grant Jensen’s lab at Caltech. The total amount of time the cells spent on ice was ∼24 h. We imaged and monitored CFUs of EG1863 cells before shipment and after 24 h of incubation on ice and observed similar growth and morphology. Upon arrival, 1 ml of the EG1863 culture was centrifuged at 3000 rpm for 5 min and resuspended in fresh PYE to a final OD_600_ of ∼8. This resuspension was mixed with fiducial markers (10 nm gold beads treated with bovine serum albumin to prevent aggregation) and 2 μl of the resuspension mixture was plunge-frozen on EM grids in a mixture of liquid ethane and propane [44]. Images were acquired using a 300 keV Polara transmission electron microscope (FEI) equipped with a GIF energy filter (Gatan) and a K2 Summit direct detector (Gatan). Tilt-series were collected from −50° to +50° in 1° increments at magnification of 22,500X using UCSF Tomography software [45] with a defocus of −12 μm and total dosage of 180 e^-^/Å^2^. Tomograms were calculated using IMOD software [46].

### Protein purification

Rosetta pLysS *E. coli* cells containing overexpression plasmids for AmiC, LdpF, DipM, and truncated LytM domain protein variants, all with a His_6_-SUMO tag fused to the N-terminus, were purified as described previously with minor changes [15]. Rosetta cells containing the constructs were grown in 1 L of LB at 30°C to an OD_600_ of 0.4 and then induced with 1 mM IPTG for 4 h. Cells were collected by centrifugation at 6000 x g at 4°C for 10 minutes and resuspended in 40 ml Column Buffer A (CBA: 50 mM Tris-HCl pH 8.0, 300 mM NaCl, 10% glycerol, 20 mM imidazole) per 1 L of culture. Cells were snap-frozen in liquid nitrogen and stored at −80°C until use. Pellets were thawed at 37°C and lysozyme was added to 1 μg/ml and MgCl_2_ to 2.5 mM. Cell suspensions were left on ice for 45 minutes, then sonicated and centrifuged for 30 minutes at 15,000 x g at 4°C. The protein supernatant was filtered and loaded onto a HisTrap FF 1ml column (GE Life Sciences) pre-equilibrated with CBA. The protein was eluted with 30% Column Buffer B (same as CBA except with 1M imidazole). The protein fractions were combined and His_6_-Ulp1 (SUMO protease) was added (1:500 Ulp1:protein molar ratio). The protease and protein fractions were dialyzed overnight at 4°C into CBA. Cleaved protein was run over the same HisTrap FF 1mL column equilibrated in CBA and the flow-through was collected. Flow-through fractions were dialyzed overnight at 4°C into Storage Buffer (50 mM HEPES-NaOH pH 7.2, 150 mM NaCl, 10% glycerol). Dialyzed protein was then concentrated (if needed), snap-frozen in liquid-nitrogen, and stored at −80°C.

### RBB labeled sacculi preparation

Sacculi were prepared from strain EG865 as described in [23]. *C. crescentus* cells were grown in 1 L of PYE at 30°C, collected at an OD_600_ of 0.6 by centrifugation at 6,000 x g for 10 minutes, and resuspended in 10 ml of PBS. The cell suspension was added drop wise to 80 ml of boiling 4% sodium dodecyl sulfate (SDS) solution. Cells were boiled and mixed for 30 minutes and then incubated overnight at room temperature. Sacculi were then pelleted by ultra-centrifugation at ∼80,000 x g for 60 minutes at 25°C. Pelleted sacculi were then washed four times with ultra-pure water and resuspended in 1 ml of PBS and 20 μl of 10 mg/ml amylase and incubated at 30°C overnight. The next day, sacculi were pelleted at ∼400,000 x g for 15 minutes at room temperature, washed three times with ultra-pure water, and resuspended in 1 ml of water. The sacculi suspension was labeled with 0.4 ml of 0.2 M remazol-brilliant blue (RBB), 0.3 ml 5 M NaOH, and 4.1 ml of water, and incubated at 30°C overnight. The labeled solution was neutralized with 0.4 ml of 5 M HCl and 0.75 ml of 10X PBS. Labeled sacculi were pelleted at 16,000 x g for 20 minutes at room temperature. The pellet was washed with water until the supernatant was clear. Blue-labelled sacculi were resuspended in 1 ml of 0.2% azide, incubated at 65°C for 3 hours, and then stored at 4°C.

### Dye-release assay

The dye release assay was adapted from [23]. Briefly, 10 μl of RBB-labeled sacculi was incubated at 30°C for 3 hours with AmiC, LdpF variants, DipM variants, or FtsX ECL singly or in combination. All proteins were used at 4 μM. Total reaction volumes were brought to 100 μl with PBS. Lysozyme (4 μM) was used as a positive control. After 3 hours of incubation, reactions were heat inactivated at 95 °C for 10 minutes and centrifuged for 20 minutes at 16,000 x g. Supernatants were collected and the absorbance was measured at OD_595_.

## Acknowledgements

We thank Jasmine Burrell for her work characterizing stalk protein localization in Δ*ftsE*; Martin Thanbichler for sharing plasmids, strains and preliminary LdpF data; Yves Brun, Erkin Kuru, and Michael van Nieuwehnze for HADA; Yves Brun and Ellen Quardokus for assistance with TEM and for sharing the StpX-GFP plasmid; and Zhuo Li for providing the wild-type stalks images in Fig. S3. This work was supported in part by the National Institutes of Health grants R35 GM122588 (to GJJ) and R01 GM108640 (to EDG).

## Supporting Information Captions

**Fig S1**. *In vivo* LdpF-mCherry has a diffuse, patchy localization in WT and Δ*ftsE*.

(A) Predicted domain organization of *E. coli* EnvC and *C. crescentus* LdpF. (B) Localization of LdpF-mCherry induced for 4 h in WT or Δ*ftsE*. Abbreviations are as follows: SS = signal sequence; CC = coiled coil domain; LytM = LytM domain. Scale bar = 2 μm.

**Fig S2**. LdpF binds the extracellular loop (ECL) of FtsX but does not activate AmiC *in vitro*.

(A) Bacterial two-hybrid of T18 and T25 fusions to the ECL of FtsX, the coiled coil domain (CC) of LdpF, and the CC cytoplasmic protein ZauP, which was used as a negative control. (B,C) Dye release assay with RBB-labeled sacculi and purified variants of AmiC, LdpF, DipM, and the ECL of FtsX. Each protein was used at 4 μM and reactions were incubated at 30^°^C for 3 h. Reactions were performed in triplicate. *** = p < 0.0001 by one-way ANOVA.

**Fig S3**. Electron cryotomography (ECT) of cells lacking FtsE and AmiC reveal stalk-like connections of heterogeneous widths.

(A) *ftsE* mutant with skinny connections that are stalk-like, but have regions with heterogeneous widths. (B) WT *C. crescentus* stalks with cross-bands. # = cross-band. Scale bar (A) = 100 nm; Scale bars (B) = 200 nm.

**Fig S4**. *ftsE* mutants incorporate new cell wall material throughout skinny connections between cell bodies.

HADA labeling of (A) WT, (B) Δ*ftsE*, and (C) Δ*ftsE* cells depleted for AmiC for 6 h. (D) FtsZ-CFP localization after 1 h of induction in Δ*ftsE* cells. * = HADA incorporation throughout skinny connections in Δ*ftsE*; # = absence of FtsZ at skinny connections in Δ*ftsE*. Scale bars = 2 μm.

**Fig S5**. Overproducing LdpF, the LytM domain (LytM) of LdpF or AmiC is not lytic *in vivo*.

Phase contrast images of cells overproducing LdpF, the LytM of LdpF or AmiC for 24 h. Scale bar = 2 μm.

**Text S1.** Supporting results and discussion describing biochemical investigation of cell wall hydrolase activities of LytM proteins and AmiC.

**Table S1.** Strains and plasmids used in this study with their methods of construction.

